# Comparative analysis of kidney organoid and adult human kidney single cell and single nucleus transcriptomes

**DOI:** 10.1101/232561

**Authors:** Haojia Wu, Kohei Uchimura, Erinn Donnelly, Yuhei Kirita, Samantha A. Morris, Benjamin D. Humphreys

## Abstract

Kidney organoids differentiated from human pluripotent stem cells hold great promise for understanding organogenesis, modeling disease and ultimately as a source of replacement tissue. Realizing the full potential of this technology will require better differentiation strategies based upon knowledge of the cellular diversity and differentiation state of all cells within these organoids. Here we analyze single cell gene expression in 45,227 cells isolated from 23 organoids differentiated using two different protocols. Both generate kidney organoids that contain a diverse range of kidney cells at differing ratios as well as non-renal cell types. We quantified the differentiation state of major organoid kidney cell types by comparing them against a 4,259 single nucleus RNA-seq dataset generated from adult human kidney, revealing immaturity of all kidney organoid cell types. We reconstructed lineage relationships during organoid differentiation through pseudotemporal ordering, and identified transcription factor networks associated with fate decisions. These results define impressive kidney organoid cell diversity, identify incomplete differentiation as a major roadblock for current directed differentiation protocols and provide a human adult kidney snRNA-seq dataset against which to benchmark future progress.

## Introduction

Chronic kidney disease affects 26 - 30 million adults in the United States and 11% of individuals with stage 3 CKD will eventually progress to end stage renal disease (ESRD) - requiring dialysis or kidney transplantation.^1^ In 2015, 18,805 kidney transplants were performed in the United States, but 83,978 patients were left waiting for a transplant due to a shortage of organs.^2^ Development of new treatments to slow progression of CKD are desperately needed but progress has been slow in part because the kidney is a complex organ and although rodent models are commonly used, it is increasingly appreciated that humans and mice are more transcriptionally different than they are similar.^3^

In this context, the emergence of methods to direct the differentiation of pluripotent human stem cells (PSC) to kidney organoids has been received with great excitement.^4-8^ Over the last three years several groups have published stepwise protocols, all based upon kidney development during embryogenesis, resulting in generation of kidney tissue in vitro.^6, 9^ These protocols modulate activity of several signaling pathways but principally the Wnt and Fgf pathways, to generate renal progenitor populations that ultimately self organize. Mature organoids contain up to hundreds of nephron structures including glomeruli, properly segmented tubules and interstitial cell types.

The ability to grow kidney organoids from patient-derived tissue offers unprecedented opportunities for the investigation of human kidney development, homeostasis and disease. For example, kidney organoids have been used to successfully model autosomal dominant polycystic kidney disease,^10^ acute kidney injury^11^ and vascularization of the glomerular tuft.^12^ A longterm goal is to generate transplantable kidneys grown in the laboratory though this is far off.^13^ However, bulk transcriptional analysis of kidney organoids suggests that they are more similar to first trimester human kidney rather than mature kidney.^4^ Further, exactly which cells are generated and their degree of maturation remain undefined. This information is required to leverage kidney organoids for investigation of the most common adult kidney diseases such as CKD, diabetic nephropathy and acute kidney injury.

Here we have used scRNA-seq and single nucleus RNA-seq (snRNA-seq) to generate the first comprehensive molecular maps describing kidney organoid cell diversity in two separate, commonly employed differentiation protocols, as well as in adult human kidney. Our analysis reveals new insights including: (1) both protocols generate at least 12 separate kidney cell types; (2) off-target nonrenal cell types are present in kidney organoids; (3) lineage relationships revealed through pseudotemporal ordering during kidney organoid differentiation; (4) kidney organoid cell types are immature when benchmarked against adult human kidney single nucleus RNA-seq. These datasets provide a novel framework for judging the success of and improving kidney differentiation protocols towards mature human kidney tissue.

## Results

### Single Cell RNA-seq Defines Cell Diversity in Kidney Organoids

We used the human induced pluripotent stem cell (iPSC) line BJFF.6, created from male foreskin fibroblasts and reprogrammed with Sendai virus. We confirmed that the BJFF.6 line could efficiently generate kidney organoids using both the protocol described by Takasato et al. (9, 13), and the protocol described by Morizane et al. (6, 8) (Figure 1A-B, hereafter referred to as the Little or Bonventre protocol, respectively). Each protocol generated nephron-like structures that closely resembled published reports (Figure 1C-F).

**Figure 1:**
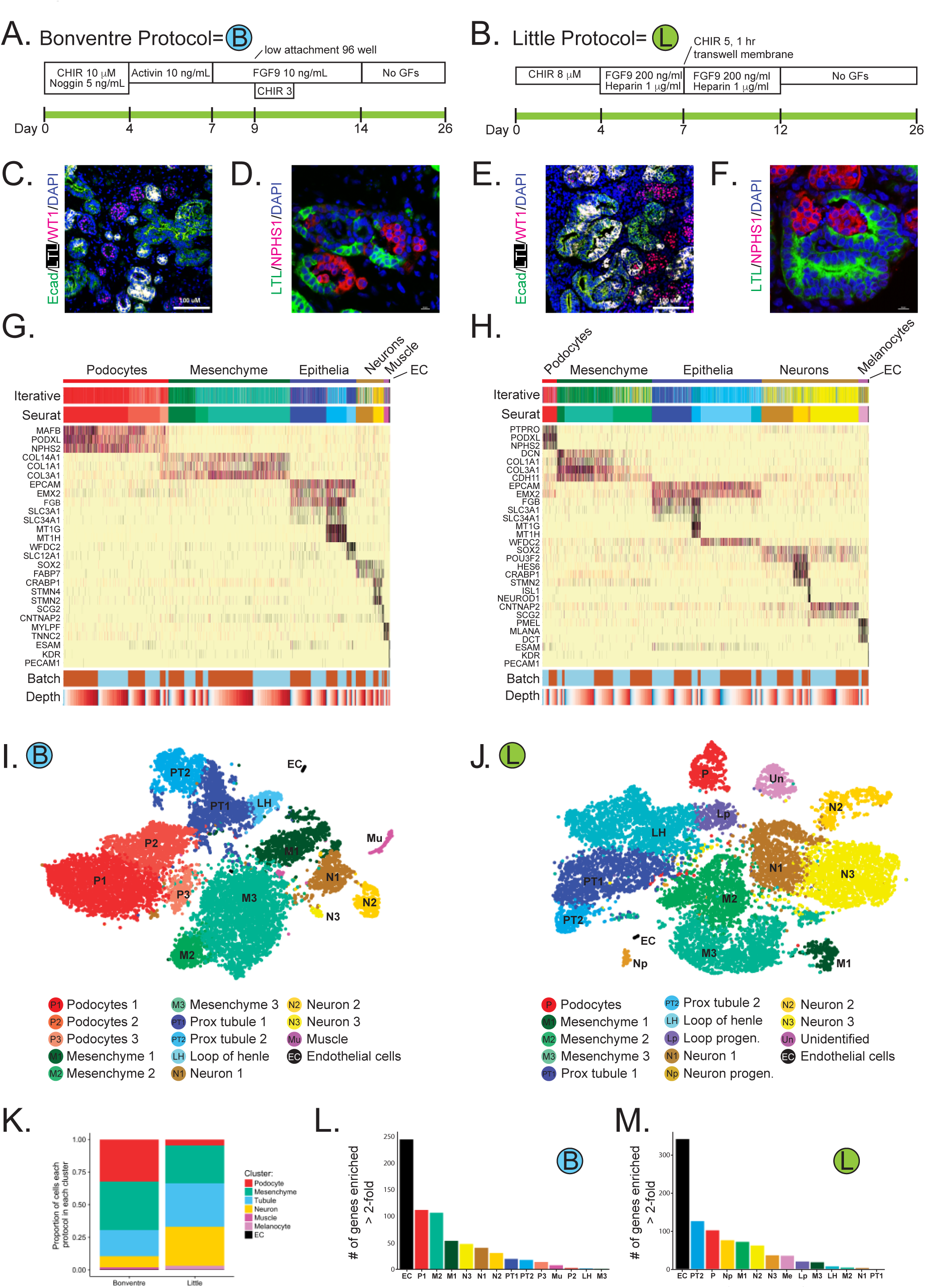
Comprehensive single-cell RNA sequencing demonstrates development of a spectrum of cell types in kidney organoids. (**A,B**) Diagram of human iPS directed differentiation protocols. (**C-F**) Immunofluorescence analysis of day 26 organoid for proximal tubule (LTL), distal tubule (ECAD), and podocytes (WT1 and NPHS) from Bonventre protocol (**C,D**) and Little protocol (**E,F**). (**G,H**) Heatmap of all cells clustered by recursive hierarchical clustering and Louvain-Jaccard clustering (Seurat), showing selected marker genes for every population of Bonventre protocol (**G**) and Little protocol (**H**). The bottom bars indicate the batch of origin (“Batch”) and number of UMI detected/cell (“Depth”). (**I,J**) tSNE plot of cells based on the expression of highly variable genes for the day 26 organoids from Bonventre protocol (**I**) and Little protocol (**J**). The detected clusters are indicated by different colors. (**K**) Comparison of the fraction of each major cell population between the two protocols. (**L,M**) Number of genes that are at least twice as highly expressed in a specific cell cluster as in the second-highest-expressing cell cluster in Bonventre protocol (**L**) and Little protocol (**M**), only including for which the differential expression between the first- and second-highest expressing clusters is significant (q < 0.05).

Using DropSeq, we isolated and sequenced mRNA from a total of 29,922 cells harvested from day 26 organoids. The Little protocol generated larger organoids, so we sequenced one each from separate batches. For the smaller Bonventre protocol organoids, we combined 8 organoids each from two separate batches. Analysis of sequencing reads revealed about ~2,200 uniquely detected transcripts from ~1250 genes for each cell (Supplementary Table 1). After correcting for batch effects, we reduced the dimensionality of the organoid data by running principal component (PC) analysis on the most highly variable genes, and then performed graph-based clustering on the significant PCs and finally visualized the distinct sub-group of cells using t-distributed stochastic neighbor embedding (tSNE, Figure 1G-J). This unsupervised clustering approach identified 14 transcriptionally distinct populations present in organoids generated from either the Bonventre or the Little protocol.

We performed a number of quality control steps. To examine potential batch effects and to quantify variability among organoids, we projected cells from different batches onto the same tSNE diagram, which showed that cells were intermixed regardless of batch (Supplementary Figure S1AB). Furthermore, cluster-based correlation analysis on both protocols revealed that the correlation for the cells in the same cell cluster from different batches was always greater than the correlation for cells in the same batch from different cell clusters (Supplementary Figure S1C,D). In addition, using an alternative clustering approach, iterative hierarchical clustering,^14^ we identified the same major organoid cell populations (Figure 1G-H). We did observe variability in the relative size of the clusters between batches particularly for the small cell clusters in Little protocol. For example, one neuron-like cluster consisting of 0.7% total cells was present only in batch 2 in the Little protocol (Figure 1K).

We assigned identities to all clusters by comparing unique transcripts expressed specifically in each cluster with existing RNA-seq datasets and the literature. The largest clusters in the Bonventre organoids were podocytes (32.2%) whereas the largest cluster in the Little protocol were tubular cells (33.5%). The smallest clusters were endothelial cells in both protocols, making up 0.3% or less of total cells. Although each protocol broadly generated similar kidney cell types, there were substantial differences in the comparative fractions of these cell types between protocols. For example, the Bonventre protocol generated three separate podocyte clusters representing embryonic and more mature podocytes whereas the Little protocol generated only one podocyte cluster. Notably, the Bonventre protocol podocyte clusters together represented 32.2% of total organoid cells, whereas Little organoid podocytes made up only 4.5% of cells.

We also observed substantial numbers of non-renal cell types in both protocols. Organoids generated using the Little protocol contained four neuronlike clusters and one cluster that we could not annotate but that expressed some melanocyte markers. Together, these non-renal cell types made up 32.8% of the total cell population at day 26 (Figure 1K). Bonventre organoids also had three neuron-like clusters, as well as a muscle-like cell cluster, comprising a total of 10.1% of the total cell count. Some clusters had relatively few genes uniquely expressed, likely reflecting different developmental stages of the same cell type. To quantitate how unique each cluster was from all the other clusters, we plotted the number of genes per cluster that were at least two times as highly expressed as in the second-highest-expressing cluster (Figure 1L-M).

We confirmed differences in relative abundance of both renal and nonrenal cell types by comparing marker gene expression for podocytes (NPHS1) and loop of henle (SLC12A1) as well as muscle (MYLPF, MYOG) and neuronal (CRABP1, MAP2) by qPCR (Figure 2A-F). To localize neuron-like cells, we performed immunostaining for CRABP11, a neuronal marker expressed in neuron-like clusters from both protocols. CRABP1 protein expression localized to spindly cells present in the interstitium (Figure 2G-H). We could identify coexpression of MAP2, a microtubule-associated protein required for neurogenesis, in many CRABP1+ interstitial cells (Figure s2), further supporting a neuronal lineage. However, we could also detect coexpression of MEIS1 – a marker of kidney stroma – in some CRABP1+ cells as well (Supplemental Figure S3A-C). Gene imputation analysis showed CRABP1 expression to be present in a small subset of mesenchymal cells from both protocols (Figure S3D,E). Since CRABP1 was recently identified as a marker of stromal progenitors in e14.5 kidney,^15^ we tested whether CRABP1 expression might mark a stromal progenitor cluster. The correlation between our neural clusters and e14.5 mouse stroma was very poor, however (Supplemental Figure S4) and very few of the top 50 stromal progenitor genes were coexpressed in the CRABP1 cluster (Supplemental Figure S5). These analyses confirm the predominant neural identity of CRABP1 expressing cells in the kidney organoid. Expression of CRABP1 and MAP2 was low at earlier timepoints but rose substantially by day 26 (Figure S6A-E). Re-analysis of the bulk RNA-seq data in Takasato et al.^4^ confirmed the presence of many neuronal genes identified by our analysis (Figure S6F), suggesting that some degree of differentiation towards neural fates may be a common outcome of current organoid differentiation protocols.

**Figure 2:**
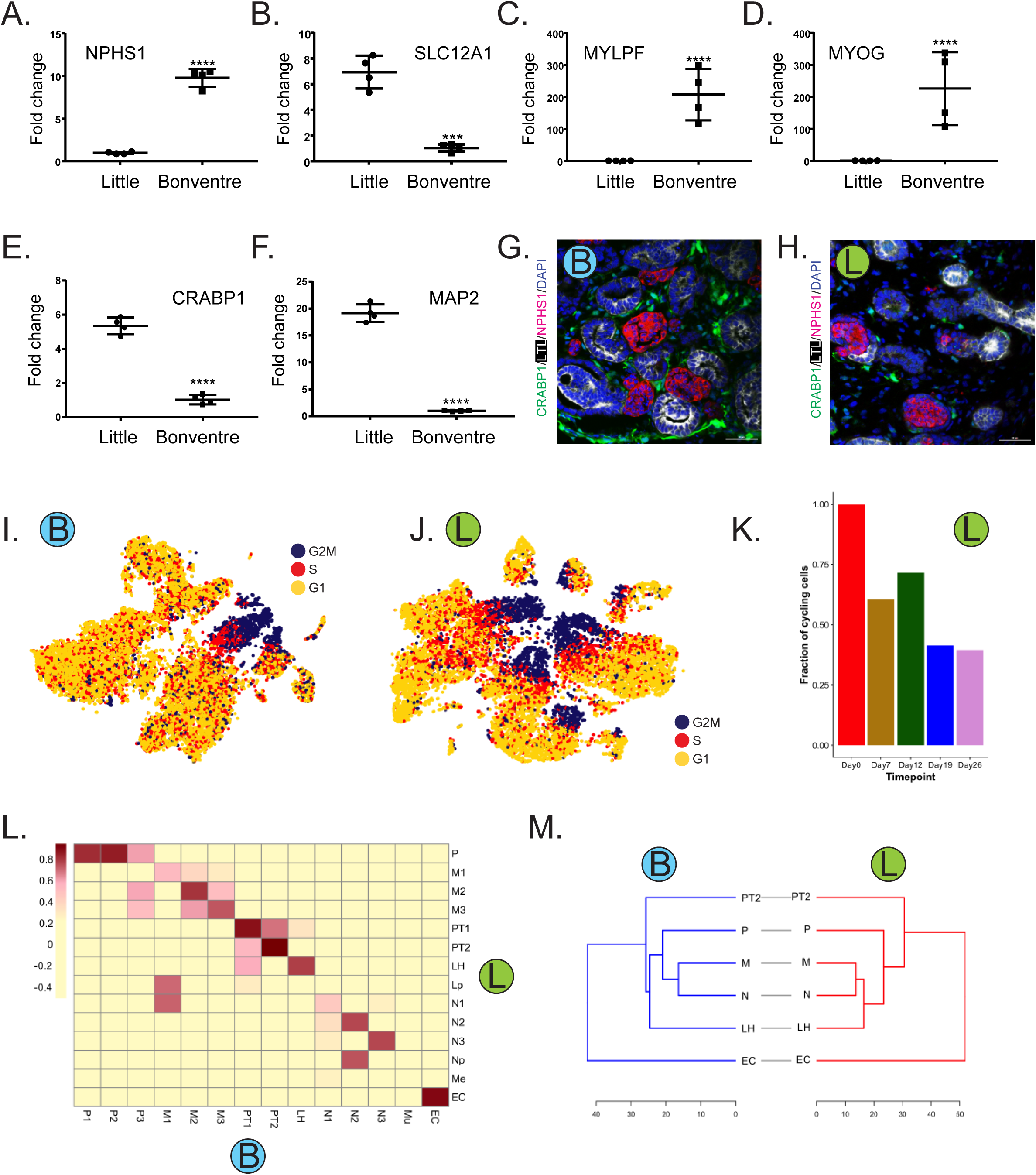
Comparison of kidney cell types, frequency and differentiation state between organoid differentiation protocols. (**A-F**) Quantitative PCR to compare cell type marker expression for podocyte (NPHS1), LOH (SLC12A1), muscle (MYLPF and MYOG), and neuron (CRABP1 and MAP2) between organoid protocols. (**G-H**) Immunofluorescence analysis of neural marker CRABP1 expression (green) in Little (**G**) and Bonventre (**H**) protocols. Cells were co-stained with PT (LTL, white) and podocyte (NPHS, red) markers. (**I-J**) tSNE plot showing cell cycle states for the organoid cells from Bonventre (**I**) and Little (**J**) protocols. (**K**) Analysis of the proliferating cells present at each time point during the organoid differentiation (Little protocol). (**L**) Heatmap indicating Pearson’s correlations on the averaged profiles among common cell types for Bonventre and Little organoids. (**H**) Dendrogram showing relationships among the cell types in Bonventre (left) and Little organoid (right). The dendrogram was computed using hierarchical clustering with average linkage on the normalized expression value of the highly variable genes.

### Cell Cycle Analysis

During kidney development, progenitor cell populations are characterized by rapid cell cycle progression whereas differentiating cell types proliferate at slower rates. We therefore analyzed cell cycle status as a proxy for degree of differentiation. We scored all cells from both protocols based on cell cycle gene expression and assigned a cell cycle phase (G2M, S or G1). The total fraction of cycling cells was similar between the protocols at 43.9% and 39.1% cycling cells in the Bonventre and Little organoids, respectively (Figure 2I,J). However, in the Bonventre organoids, rapidly cycling cells were limited to two cell clusters – a mesenchymal and neuronal cluster. By contrast, in Little organoids, rapidly cycling cells were present as a subset of 6 separate clusters (two mesenchymal, two neuron-like, one unidentified and an epithelial cluster). We interpreted the broader proliferative distribution of the Little organoid to potentially reflect that the organoid had been harvested before it was fully differentiated. Indeed, cell cycle analysis on the cells collected from different time points using Little protocol revealed that the proportion of cycling cells decreased along the kidney organoid differentiation process (Figure 2K), suggesting that the degree of cell differentiation or cell type maturity might be negatively correlated with the proliferative capacity.

### Consistent Kidney Cell Types Generated by each Protocol

We compared cell types generated across protocols by computing Pearson’s correlations on the averaged expression profiles among day 26 clusters. We found that the most highly correlated cell types were podocytes, mesenchyme, proximal tubule, endothelial cells as well as neural-like cells. We also computed a dendrogram by hierarchical clustering which revealed a deep branch separating endothelial cells from all other cell types. Mesenchyme and neural clusters were closely related in both protocols. In fact all major cell types generated by each cluster correlate very well with each other despite being generated by different protocols (Figure 2L,M).

### Kidney Organoid Cell Subsets

Re-clustering analysis of the tubular cells identified additional cell clusters in both protocols. We detected 8 and 5 tubular subtypes in Bonventre and Little organoids, respectively (Figure 3A-D). This includes a subpopulation that expressed the ureteric bud marker GATA3 in both protocols. Prior reports have suggested that kidney organoids contain derivatives of both major progenitor populations, the metanephric mesenchyme and the ureteric bud. However, the GATA3 cell cluster did not express mature collecting duct markers (e.g. AQP2, AQP4).

We next compared a panel of marker genes across podocyte, proximal tubule and loop of henle cell clusters across protocols. This revealed higher expression of podocyte developmental markers CCND1, CDH6, EMX2 and SOX4 in the Little organoids. Bonventre podocytes had higher expression of podocyte differentiation markers and lower expression of proliferation markers (Figure 3E). By contrast, for proximal tubule, Little organoids had higher expression of both differentiation markers and select developmental genes, whereas Bonventre proximal tubule had increased expression of genes that were difficult to interpret, including metal binding genes MT1M and MT1H (Figure 3F). Loop of henle was more differentiated in the Bonventre organoids (Figure 3G).

**Figure 3:**
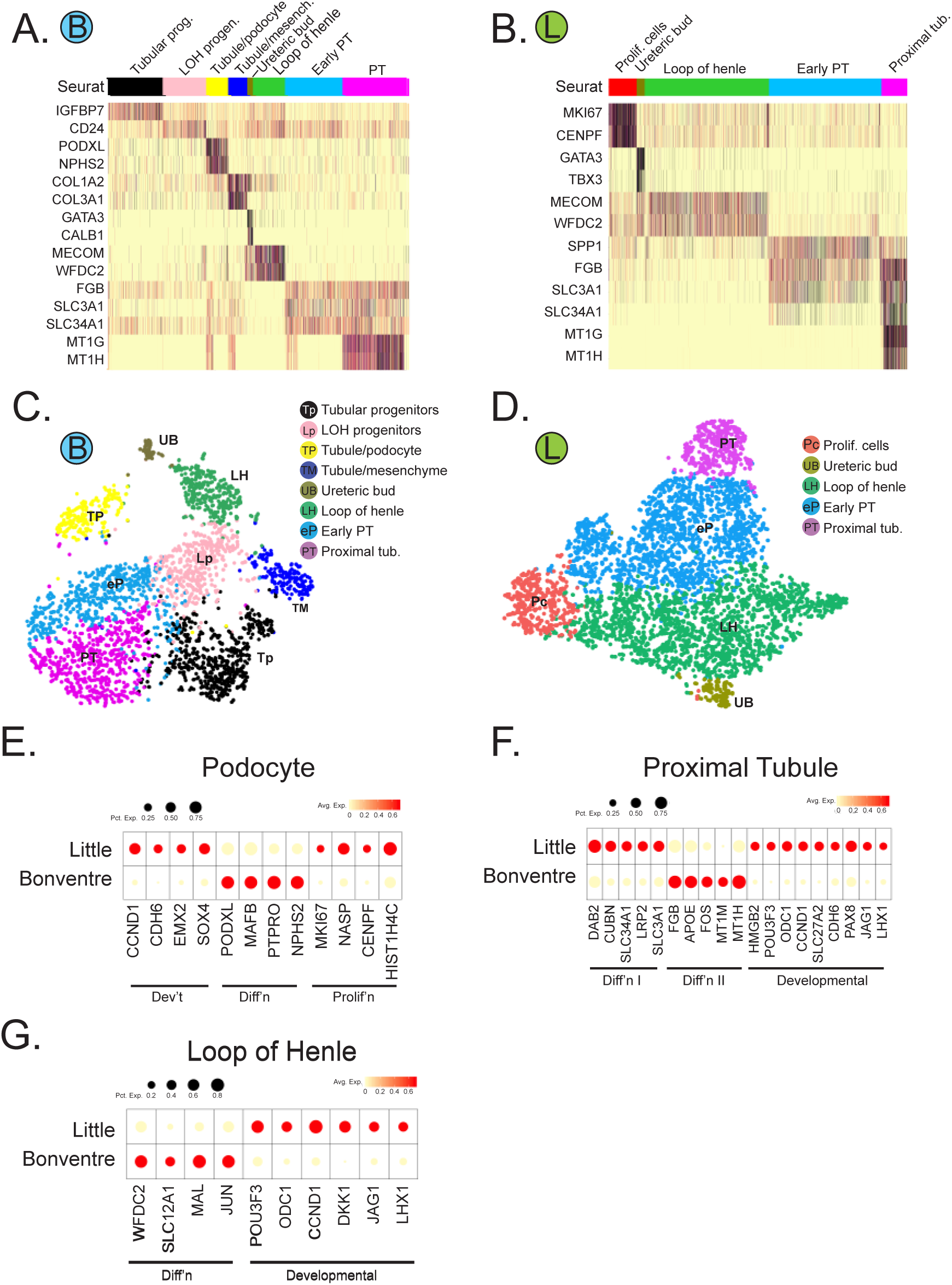
Human kidney organoids contain subclasses of tubular epithelial cells. (**A,B**) Heatmap showing selected marker genes for every tubular subpopulation of Bonventre protocol (**A**) and Little protocol (**B**). (**C,D**) tSNE plot of tubular subclusters in kidney organoid from Bonventre protocol (**C**) and Little protocol (**D**). The detected clusters are indicated by different colors. (**E-G**) Dotplot comparing the expression of cell type signature and developmental/proliferating genes on podocytes (**E**), proximal tubule (**F**), and LOH (**G**) between the two protocols.

### Organoid Kidney Cell Types are Immature

A critical question is the degree to which kidney organoid cell types resemble their native counterparts in molecular terms. We addressed this question in two ways. First, we compared organoid cell type gene signatures with a recent mouse P1 single-cell RNA-seq dataset.^16^ Despite the species differences, we detected a strong correlation between corresponding cell types for podocytes, proximal tubular cells, loop of henle, stroma and endothelial cells (Figure S7). Loop of henle cells were also correlated, albeit to a lesser extent, with distal tubule, collecting duct, and ureteric bud tips from embryonic kidney, suggesting their immaturity. For both organoid protocols, the M1 mesenchymal clusters showed medium correlation to cap mesenchyme in addition to stroma. Notably, none of the off target clusters (neural, muscle, melanocyte-like) correlated to cell types found in P1 kidney.

We next compared kidney organoid cell types with their adult human counterparts. Multiple attempts at single cell RNA-seq failed (data not shown), however we were successful in generating adult human kidney single nucleus RNA-seq data using the InDrops platform.^17^ We sequenced 4,259 nuclei to a similar depth (Table 1) as the organoid datasets, and identified 6 distinct epithelial cell clusters, including podocytes, proximal tubule, loop of henle, distal tubule, principal cells and intercalated cells (Figure 4A-C). The absence of stromal or leukocyte populations presumably reflects either dissociation bias and/or a cell frequency below our limit of detection. Single cell and single nucleus RNA-seq datasets have been shown to generate comparable results^18^ so we next correlated all kidney organoid epithelial cell types to their corresponding endogenous counterpart from human adult kidney. We observed an expected correlation between corresponding cell types of organoid and human kidney. Once again, organoid loop of henle correlated with adult loop of henle, but also distal tubule and collecting duct – likely reflecting their developmental immaturity (Figure 4D).

**Figure 4:**
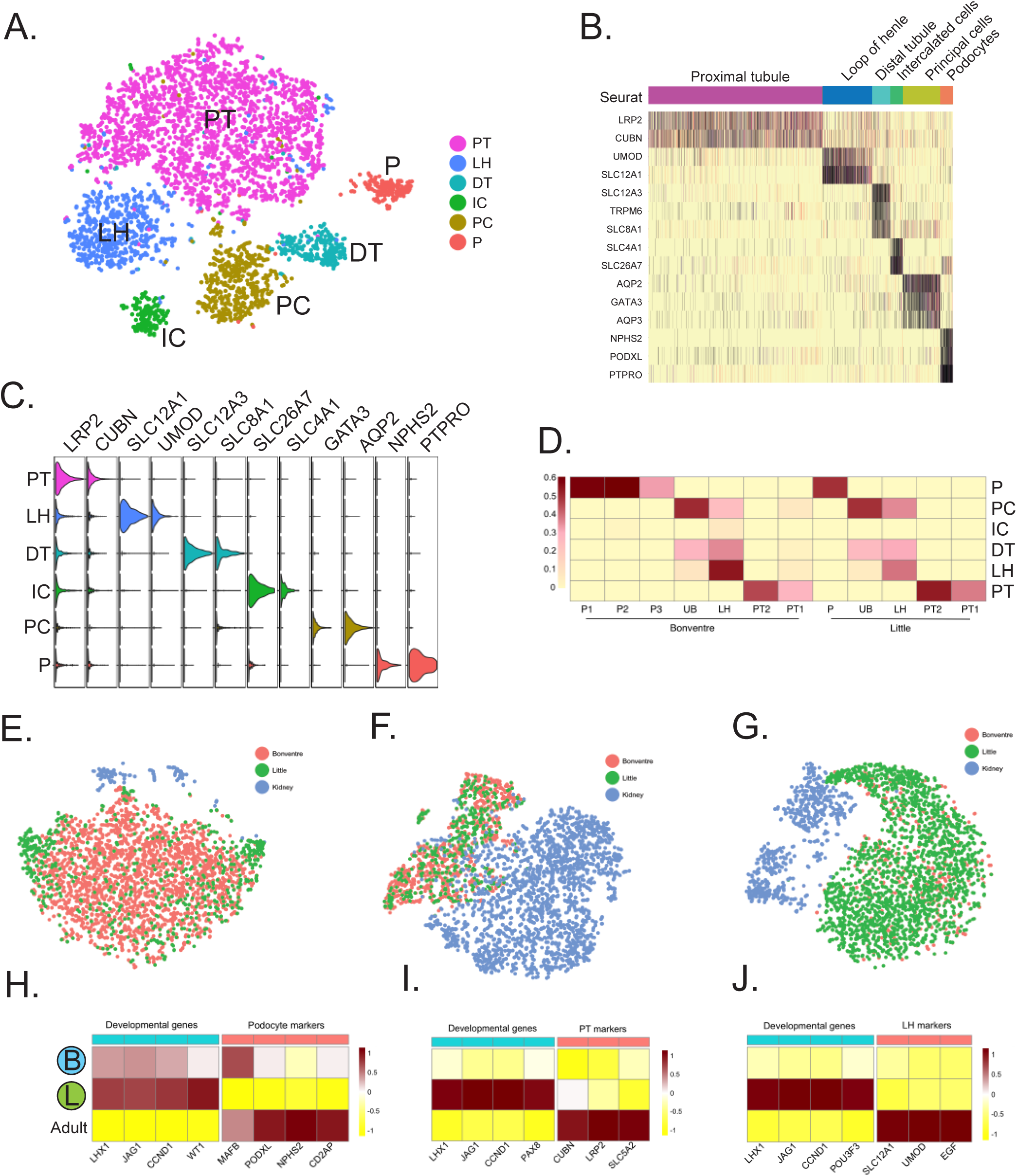
Organoid cell types are immature compared to benchmarked adult kidney cell types. (**A**) Unsupervised clustering identified six distinct cell types in human adult kidney. That includes three tubular cell types (PT, LH and DT), two collecting duct cell populations (PC and IC) and one podocyte population (P). (**B**) Heatmap showed that putative molecular signature marks the identity of each cluster. (**C**) Violin plot further confirmed the clean expression of well-known cell type specific markers in each cell population, which makes it suitable for use in benchmarked comparison analysis. (**D**) Pearson correlation analysis to compare the organoid cell types and their endogenous counterparts in human kidney. Color bar indicates the correlation score. (**E-G**) Clustering analysis on each organoid cell type and their kidney counterpart. Cell origins were visualized in tSNE. (**H-J**) Comparison of the average expression of marker genes and developmental genes between organoid cell types and adult kidney cell types. Expression value was scaled by z-score.

Although organoid and adult kidney epithelial cell types correlated well, our prior analysis suggested that organoid derived cells expressed developmental markers. To visualize these differences, we projected organoid-derived podocyte, proximal tubule and loop of henle clusters with their endogenous counterparts by tSNE (Figure 4E-G). In all cases the organoid-derived cells clustered together, but did not cluster with adult kidney clusters. Further emphasizing the substantial transcriptional differences between organoid-derived cells and adult kidney, developmental markers present in organoid cells showed much lower average expression in adult kidney. By contrast, differentiation markers for each cell type had much higher expression in adult kidney than in organoid-derived cells (Figure 4H-J).

Because transcription factors regulate cell state, we next tested the hypothesis that organoid cell immaturity might reflect partial expression of the gene regulatory network present in mature kidney cells. We found 44 transcription factors present in adult human proximal tubule, many of which have not been reported previously (Table S2). For example, High mobility group nucleosome-binding domain-containing protein 3 (HMGN3) is a thyroid hormone binding receptor that regulates gene expression, is strongly expressed in proximal tubule, and thyroid hormone is known to regulate renal fluid and electrolyte handling, suggesting HMGN3 may mediate thyroid hormone actions in the proximal tubule. In human adult podocytes, we identified 21 transcription factors, 8 of which have not been reported previously. We validated expression of six of these novel transcription factors at the protein level (Supplemental Figure S8).

Both proximal tubule cells and podocytes derived from organoids expressed only a small fraction of the transcription factors we identified in the adult cell types. For example, Little protocol proximal tubule expressed 7/44 transcription factors, and Bonventre proximal tubule only 3/44. Similarly, Little protocol podocytes expressed 8/21 transcription factors and Bonventre podocytes 9/21 compared to adult podocytes (Table S3). This result suggests that organoid cells, despite expressing some markers of differentiated cells, are fundamentally different from their terminally differentiated adult counterparts.

To test whether longer organoid incubation times might improve cell differentiation status, we grew organoids from both protocols out to 34 days and performed scRNA-seq on a total of 6,115 cells (Table 1). We compared expression of differentiation markers across clusters, and discovered that differentiation was generally worse not better at this later timepoint, with loss of endothelial cells, reduced expression of differentiation markers across most clusters, and the emergence of a novel off-target muscle cell cluster (Supplemental Figure S9).

### Disease-related genes predicted by GWAS are expressed in single cell types in adult and organoid kidney

Human kidney organoids are already being used to model monogenic human kidney diseases. However there are many more complex trait disease genes that have been identified by Genome-Wide Association Studies (GWAS). Recently Park et al. reported that many human monogenic and complex trait genes are expressed predominantly in a single mouse kidney cell type.^19^ Since human and mouse differ transcriptionally,^3^ we mapped complex trait genes associated with CKD, hypertension or plasma metabolite levels to both adult human kidney epithelial cell types as well as to kidney organoid-derived cell types. We generated GWAS gene lists including 117 genes for chronic kidney diseases, 275 genes for hypertension and 777 genes for plasma metabolite levels.

We could map expression of 143 of these genes to adult kidney epithelial cell types. We mapped 63 of these genes to Bonventre organoid cell types and 89 genes to Little organoid cell types. In all cases, we confirmed that the majority are expressed in only a single kidney cell type (Figure 5, Supplemental Figure S10). Unexpectedly, podocytes were the human kidney cell type with the highest number of hypertension genes expressed. These cells are not widely believed to play important roles in regulating blood pressure. Consistent with their central role in secretion and reabsorption, proximal tubule had the highest number of genes associated with plasma metabolite levels. There were about half as many genes that could be mapped to organoid cell types, but the pattern of single cell type expression was preserved. These results confirm and extend those of Park et al., which was performed in mouse and not human, and also suggest that kidney organoids may represent an attractive tool to begin defining the biology underlying complex trait gene associations.

**Figure 5:**
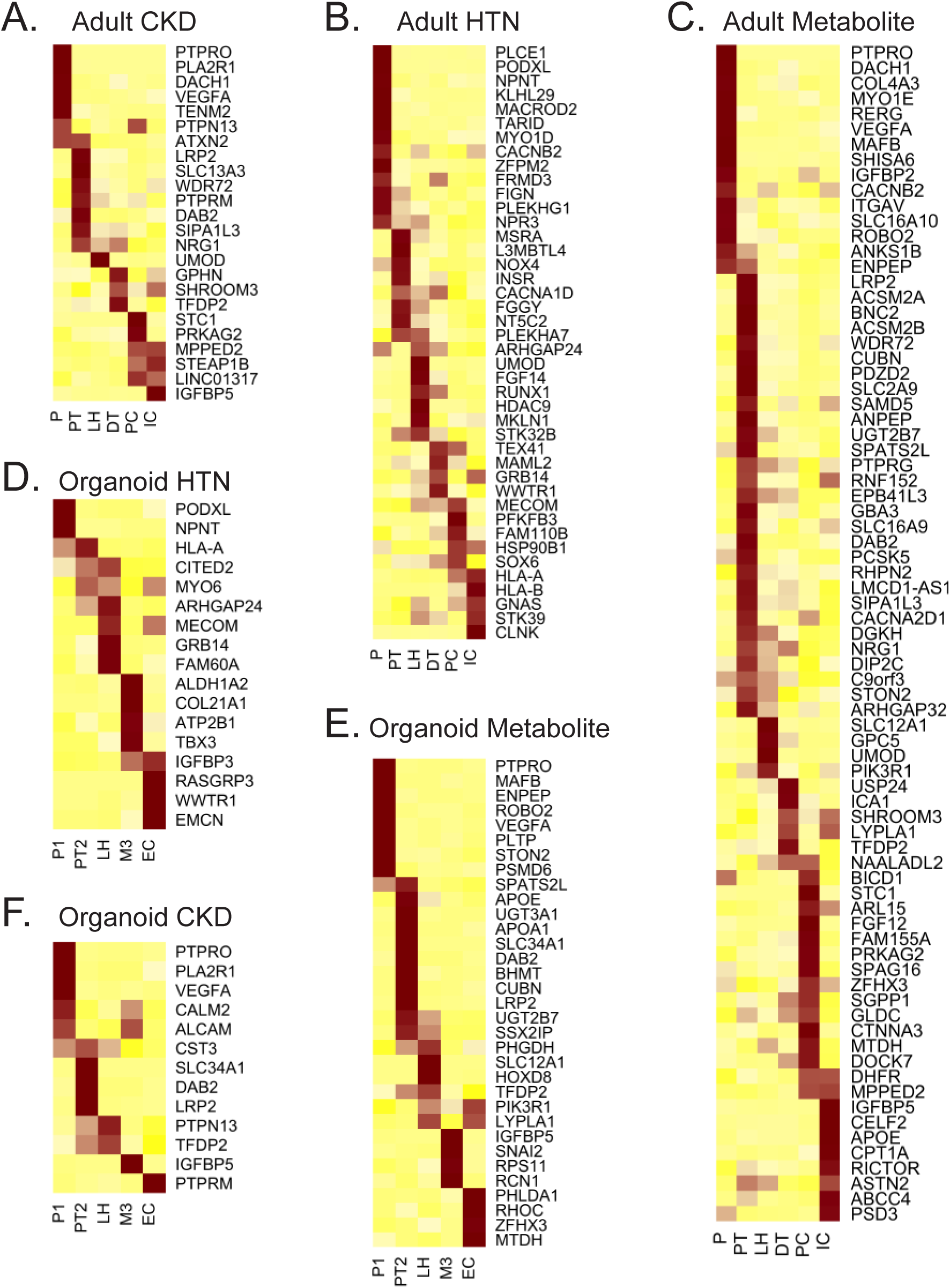
Assessing cell type specificity of genes identified in kidney disease related GWAS. Cell type specificity for genes reported in CKD related GWAS (**A-C**), hypertension related GWAS (**D-E**) and plasma metabolite levels related GWAS (**G-I**). Each gene reported in a kidney disease related GWAS was assigned to the organoid and adult kidney cell type in which it was found to be differentially expressed in each cell type (likelihood ratio test). Heatmap was used to visualize the z-score normalized average gene expression of the candidate genes for each cell cluster identified from Bonventre organoid, Little organoid and adult human kidney.

### Lineage Reconstruction during Kidney Differentiation

To explore lineage relationships and the mechanisms of cell fate decisions during kidney organoid differentiation, we performed scRNA-seq at separate timepoints during differentiation using the Little protocol (days 0, 7, 12, 19 and 26). A total of 9,190 cells from all five timepoints were projected by tSNE, and days 0, 7 and 12 each formed single distinct clusters (Figure 6). Pluripotency gene expression (e.g., POU5F1/Oct4) was completely downregulated by day 7 with upregulation of metanephric mesenchyme markers (SALL1, FGF18 and HOXB9, Figure 6B). The day 12 cluster most closely resembled the pretubular aggregate, with genes such as JAG1 and LHX1 strongly enriched at this timepoint. Multiple clusters corresponding to differentiating cell types were present at days 19 and 26, and most later clusters contained cells from both time points, reflecting asynchronous differentiation.

**Figure 6:**
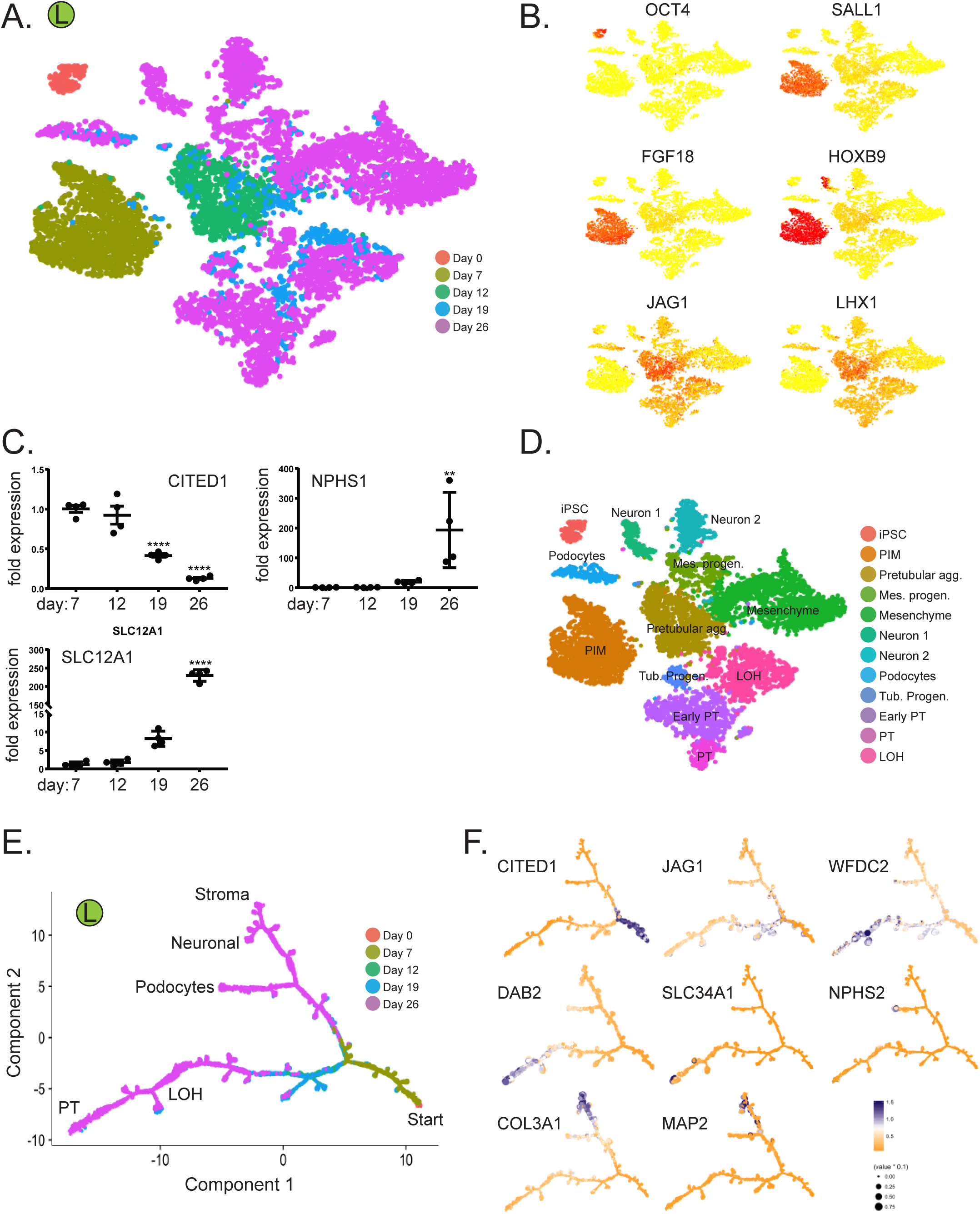
Time-course analysis of cells during organoid differentiation reveals lineage relationships. (**A,B**) Projecting cells across time-points to the tSNE. Cells were colored by the time point where they were collected (**A**) or gene expression of stage specific markers (**B**). (**C**) Validation of the stage specific marker by qPCR. \*\**p*<0.01 and \*\*\*\**p*<0.001 versus day 7. (**D**) Annotation of cell clusters based on gene expression of cell type specific markers. (**E**,**F**) Ordering of scRNA-seq expression data according to the pseudotemporal position along the lineage revealed a continuum of gene expression changes from iPSCs to differentiated cell types.

Although differentiation marker expression increased with time, and certain progenitor markers such as CITED1 decreased over time (Figure 6C), many genes marking developmental cell types persisted at day 26. Genes reflecting metanephric mesenchyme (SIX1), renal vesicle (DKK1), S-shaped body (JAG1, CCND1, CDH6 and LHX1) continued to be expressed, for example, suggesting an ongoing nephrogenic program (Figure 3E-G). Future enhancements to kidney organoid differentiation protocols will need to push maturation of these developmental states towards fully mature kidney cell types.

To detect gene expression changes during organoid differentiation, we reconstructed kidney lineage relationships by performing pseudotemporal ordering using Monocle2.^20^ The resulting cell trajectories revealed one major branch point, separating loop of henle and proximal tubule cell fates from podocyte, stromal and neural cell fates (Figure 6E). A second branch point distinguished podocyte from stromal and neural fates. Cell fates were defined by projecting marker gene expression onto the pseudotime trajectories (Figure 6F).

To explore the mechanisms underlying the first major developmental fate decision tubular epithelium fates from podocyte, stromal and neural fates, we scrutinized transcription factors expressed during this first branch point, corresponding to day 12. Using BEAM analysis from Monocle2, we found strong downregulation of HOXC6 and CDX2 just before the branch, with strong upregulation of a core set of transcription factors including EMX2, POU3F3, HES1, PAX2 and SIM1 exclusively in the tubular epithelium branch. By contrast, we noted strong upregulation of SNAI2, ETS1 and HOXC10 in the podocyte/mesenchyme branch (Supplemental Figure S11AB).

## Discussion

Fulfillment of the promise of human kidney organoids requires comprehensive characterization of their cell composition, comparison of differing protocols and a better understanding of the degree to which they produce mature, differentiated kidney cell types. Using scRNA-seq, we have established that current protocols generate a remarkable diversity of kidney cell types. We also provide the first direct comparison of separate differentiation protocols, revealing broadly similar outcomes, but important differences in cell ratio and differentiation state.

By comparing organoid derived kidney cell types with adult human kidney, we describe how epithelial cell clusters are similar or different. Although organoid-derived epithelia are already known to express an array of markers present on terminally differentiated kidney cells, our comprehensive transcriptional analysis suggests that all organoid-derived cells are substantially immature. We have also identified various off-target cell populations present in kidney organoids, primarily neural lineage cell types. Unexpectedly longer organoid incubation did not improve differentiation, but caused reduced expression of terminal markers and generated new off-target cells, suggesting kidney cell type dedifferentiation with time. These results indicate a need to identify conditions that will better support continued organoid maturation.

Clues for future improvement of differentiation protocols are suggested by the results of pseudotemporal ordering, which provides an initial roadmap to help understand lineage relationships between progenitors and more mature cell types. We envision that analysis of signaling pathways and transcription factors expressed before and after branch points will suggest potential strategies to increase or decrease specific fates in a rational fashion, for example by eliminating neural differentiation.

## Acknowledgements

This work was supported by NIH/NIDDK DK103740, by an Established Investigator Award of the American Heart Association and by Grant 173970 from the Chan Zuckerberg Initiative (all to B.D.H). KU was supported by JSPS Postdoctoral Fellowships for Research Abroad.

## Methods

No statistical methods were used to predetermine sample size. The experiments were not randomized. The investigators were not blinded to allocation during experiments and outcome assessment.

### iPSC Culture

All experiments utilized the BJFF6 human iPSC line reprogrammed by Sendai virus from human foreskin fibroblasts (Washington University Genome Engineering and iPSC Core). This line is confirmed to be karyotypically normal. BJFF6 cells were maintained in 6-well plates coated with matrigel (Corning) in Essential 8 medium (Thermo Fisher Scientific). iPSC cells were dissociated using ReLeSR (STEMCELL Technologies), confirmed to be mycoplasma free and maintained below passage 50.

### Kidney Organoid Differentiation

Kidney organoids were generated following either the protocol described by Takasato et al.^9^ or that of Morizane et al.,^6^ with minimal modifications. Briefly, for the Takasato approach, BJFF cells were treated with CHIR (8 uM, Tocris Bioscience) in basal medium - APEL 2 (STEMCELL Technologies) supplemented with 5% Protein Free Hybridoma Medium II (PFHMII, Gibco) - for 4 days, followed by FGF9 (200 ng/mL, R&D Systems) and heparin (1 ug/mL, Sigma-Aldrich) for another 3 days. At day 7, cells were collected and dissociated into single cells using 0.25 % Trypsin-EDTA (Thermo Fisher Scientific). Cells were spun down at 400 g for 3 min to form a pellet and transferred onto a transwell membrane. Pellets were incubated with CHIR (5 uM) for 1 hour and then cultured with FGF9 (200 ng/mL) and heparin (1 ug/mL) for 5 days. For the next 13 days organoids were cultured basal medium changed every other day. For the Morizane approach, BJFF cells were treated with CHIR (10 uM) and Noggin (5 ng/mL, PeproTech) in basal medium - Advanced RPMI 1640 medium (Gibco) supplemented with 1X L-GlutaMAX (Thermo Fisher Scientific) - for 4 days, followed by 3 days Activin (10 ng/mL, R&D Systems) and 2 days FGF9 (10 ng/mL) treatment. At day 9, the cells were dissociated with Accutase (StemCell technologies) and resuspended in the basic differentiation medium with CHIR (3 μM) and FGF9 (10 ng/mL), and placed in ultra-low attachment 96-well plates. Two days later the medium was changed to basal medium containing FGF9 (10 ng/mL) and cultured for 3 more days. After that, the organoids were cultured in basal medium with no additional factors until harvest at day 26.

### DropSeq single-cell RNA sequencing

Organoids were dissociated using TrypLE Select (Thermo Fisher Scientific) at 37°C with shaking for 5 min and counted by hemocytometer (INCYTO C-chip) and resuspended in PBS + 0.01% BSA. Single cells were coencapsulated in droplets with barcoded beads exactly as described.^21^ Libraries were sequenced on a HiSeq 2500. We routinely tested our DropSeq setup by running species mixing experiments prior to running on actual sample to assure that the cell doublet rate was below 5%.

### Sample Preparation and inDrop single-nucleus RNA-seq of human kidney

Institutional review board approval for research use of human tissue was obtained from Washington University. Samples from a discarded human donor kidney was obtained and donor anonymity preserved. The donor was a 70 year-old white male with a serum creatinine of 1.1 mg/dL. Nuclei were isolated with Nuclei EZ Lysis buffer (Sigma #NUC-101) supplemented with protease inhibitor (Roche #5892791001) and RNase inhibitor (Promega #N2615, Life Technologies #AM2696). Samples were cut into <2 mm pieces and homogenized using a Dounce homogenizer (Kimble Chase #885302-0002) in 2ml of ice-cold Nuclei EZ Lysis buffer and incubated on ice for 5 min with an additional 2ml of lysis buffer. The homogenate was filtered through a 40-μm cell strainer (pluriSelect #4350040-51) and then centrifuged at 500 × for 5 min at 4 °C. The pellet was resuspended and washed with 4 ml of the buffer and incubated on ice for 5 min. After another centrifugation, the pellet was resuspended with Nuclei Suspension Buffer (1× PBS, 0.07% BSA, 0.1% RNase inhibitor), filtered through a 20μm cell strainer (pluriSelect 43-50020-50) and counted. RNA from single nucleus was encapsulated, barcoded and reversed transcribed using an InDrop microfluidics system (1CellBio). The library was sequenced in HiSeq2500 with custom primers.

### Immunofluorescence

Organoids were fixed in 4% paraformaldehyde (Electron Microscopy Services), cryoprotected in 30% sucrose solution overnight and embedded in optimum cutting temperature (OCT) compound (Tissue Tek). Organoids were cryosectioned at 7μm thickness and mounted on Superfrost slides (Thermo Fisher Scientific). Sections were washed with PBS (3 times, 5 minutes each), then blocked with 10% normal goat serum (Vector Labs), permeabilized with 0.2% Triton-X100 in PBS and then stained with primary antibody specific for mouse anti-WT1 (1:200, Santa Cruz Biothechnology, #SC-7385), rat anti-ECAD (1:200, Abcam, #ab11512), biotinylated LTL (1:200, Vector Labs, #B-1325), sheep anti-NPHS1 (1:200, R&D Systems, #AF4269) and rabbit anti-CRABP1 (1:200, Cell Signaling, #13163), chicken anti-MAP2 (1:200, Abcam, #ab5392) and mouse anti MEIS1 (1:200, Active Motif, #39795). Secondary antibodies included FITC-, Cy3, or Cy5-conjugated (Jackson ImmunoResearch). Then, sections were stained with DAPI (4’,6’-diamidino-2-phenylindole) and mounted in Prolong Gold (Life Technologies). Images were obtained by confocal microscopy (Nikon C2+ Eclipse; Nikon, Melville, NY).

### Statistical analysis

Data were presented as mean ± SEM. ANOVA with post hoc Bonferroni correction was used for multiple group comparison. Student t-test was used to compare 2 different groups. Graph-Pad Prism software, version 6.0c (GraphPad Software Inc., San Diego, CA) and SPSS version 22 were used for statistical analysis. P-value < 0.05 was considered as statistical significant difference.

